# Comprehensive comparison between azacytidine and decitabine treatment in an acute myeloid leukemia cell line

**DOI:** 10.1101/2022.02.03.476906

**Authors:** Tina Aumer, Constanze B. Gremmelmaier, Leander S. Runtsch, Nur Yeşiltaç, Stefanie Kaiser, Franziska R. Traube

## Abstract

Azacytidine (AzaC) and decitabine (AzadC) are cytosine analogs that covalently trap DNA methyltransferases, which place the important epigenetic mark 5-methyl-2’-deoxycytidine by methylating 2’-deoxycytidine (dC) at the C5 position. AzaC and AzadC are used in the clinic as antimetabolites to treat myelodysplastic syndrome and acute myeloid leukemia and are explored against other types of cancer. Although their principal mechanism of action is known, the downstream effects of AzaC and AzadC treatment are not well understood and the cellular prerequisites that determine sensitivity towards AzaC and AzadC remain elusive. Here, we investigated the effects and phenotype of AzaC and AzadC exposure on the acute myeloid leukemia cell line MOLM-13. We found that while AzaC and AzadC share many effects on the cellular level, including decreased global DNA methylation, increased formation of DNA double strand breaks, transcriptional downregulation of important oncogenes and similar changes on the proteome level, AzaC failed in contrast to AzadC to induce apoptosis in MOLM-13. The only cellular marker that correlated with this clear phenotypical outcome was the level of hydroxy-methyl-dC, an additional epigenetic mark that is placed by TET enzymes and repressed in cancer cells. Whereas AzadC increased hmdC substantially in MOLM-13, AzaC treatment did not result in any increase at all. This suggests that hmdC levels in cancer cells should be monitored as a response towards AzaC and AzadC and considered as a biomarker to judge whether AzaC or AzadC treatment leads to cell death in leukemic cells.

## Introduction

5-Azacytidine (azacytidine, AzaC) and 5-aza-2’-deoxycytidine (decitabine, AzadC) are cytosine analogs that belong to the compound class of hypomethylating agents and are applied in the clinic against myelodysplastic syndrome (MDS) and acute myeloid leukemia (AML) [1, 2]. Furthermore, there is ongoing research if and how AzaC and AzadC can contribute to the treatment of other types of cancer [3-5]. AzaC and AzadC have a dual mode of action by addressing epigenetic and DNA damage processes. After uptake, the majority of AzaC is incorporated into RNA, where it inhibits the tRNA (cytosine(38)-C(5))-methyltransferase (*TRDMT1*, DNMT2). In addition, it is metabolized on the diphosphate level to the respective AzadC analogue and is subsequently incorporated into DNA. After genomic incorporation, DNA methyltransferases (DNMTs) are inhibited by creating permanent crosslinks between the protein and the 5-aza-cytosine nucleobase [6, 7]. On the epigenetic level, this leads to a global loss of the epigenetic mark 5-methyl-2’-deoxycytidine (mdC) as newly synthesized DNA cannot be methylated anymore [1, 8]. Aberrant mdC patterns are one characteristic of many cancer types since mdC in promoter regions results in gene silencing [9, 10]. The induction of massive DNA demethylation in promoter regions of previously silenced tumor suppressor genes might therefore be a mean of clinically reactivating them [11]. DNA-protein crosslinks on the other hand are a severe form of DNA damage that must be repaired *via* Fanconi anemia-dependent homologous recombination (FA pathway) [12]. Unrepaired crosslinks lead to a stalled replication fork that eventually collapses and results in DNA double strand breaks (DSBs) that are highly deleterious for untransformed as well as cancer cells and therefore trigger apoptosis [13, 14]. Although the basic chemistry of AzaC and AzadC is known, many aspects of their mode of action are not well understood. It remains unclear, how much the incorporation of AzaC into RNA and thus inhibition of DNMT2 and other m^5^C methyltransferases in general contribute to the efficacy of AzaC [8]. While reports show that in some cancer cell lines AzaC is more efficient than AzadC in inducing cell death, other cancer cell lines show the opposite behavior [6, 15]. Even though the direct effects on the mdC levels have been intensively studied, there is a lack of information on how AzaC and AzadC treatment affect the proteome of AML cells. Moreover, not a single reliable biomarker has been found that can predict whether treatment with AzaC or AzadC is effective [16, 17]. Therefore, to fully exploit the potential of 5-aza-cytosines as epigenetic drugs for cancer therapy, it is of great importance to gain deeper knowledge of the molecular patterns that determine which cellular signaling pathways are altered after treatment with AzaC or AzadC.

## Results

### Effect of AzaC and AzadC on mdC, m^5^C and DNA damage in MOLM-13

To compare the effects of AzaC and AzadC treatment, we performed a comprehensive analysis on the cellular phenotype and the underlying molecular changes between AzaC- and AzadC-treated cells. As a model cell line, we chose the AML cell line MOLM-13 because previous studies reported an effective response to AzaC-as well as AzadC-treatment on DNA methylation level [7, 18], confirming that both compounds are taken up and metabolized correctly. First, we checked by ultra-HPLC triple quadrupole mass spectrometry (UHPLC-QQQ-MS) whether the global mdC level changed upon AzaC and AzadC treatment as expected. In accordance with previously published results [7], mdC levels decreased after exposure to AzaC and AzadC for 72 h (Fig. 1a). For AzaC, the decrease was concentration dependent with 0.5 µM of AzaC being insufficient to reduce mdC levels significantly compared to the control. In contrast, mdC levels dropped substantially after exposure to 0.5 µM, 1.0 µM and 2.5 µM of AzadC, without difference between the concentrations. When comparing the change of mdC levels after exposure to equal amounts of the two compounds, only the highest dose of AzaC (2.5 µM) reduced the mdC levels to a similar extent as AzadC (Fig. 1a). Next, we measured the amount of 5-methyl-cytidine (m^5^C) per tRNA (Fig. 1b). m^5^C is the product of DNMT2, which methylates cytidine 38 in tRNA^Asp^ [19] and NSUN2, which methylates various tRNAs at position 34 and 48 – 50 [20]. We observed the anticipated reduction of m^5^C in tRNA after exposure to AzaC, but not after AzadC treatment. Afterwards, we assessed the level of DNA damage that is introduced by AzaC and AzadC. Therefore, we investigated whether AzaC and AzadC treatment lead to DSB formation in a dose dependent manner in MOLM-13 by immunoblot analysis against *γ*H2AX (Fig. 1c, Figure S1a). H2AX is a histone variant that is placed at DSB sites and subsequently phosphorylated at Ser-139 (*γ*H2AX) to mark the site where the strand breaks occurred and subsequently recruit the repair machinery [21]. Since one *γ*H2AX is placed per DSB, it is a very sensitive and quantitative marker for DSB formation. For AzadC, we observed a substantial increase in *γ*H2AX levels for all three concentrations tested, whereby treatment with 2.5 µM AzadC seemed to result in slightly more DSBs than treatment with 0.5 µM and 1.0 µM. Treatment with AzaC also lead to DSB formation, although to a lesser extent than AzadC treatment and in a strictly concentration-dependent manner.

**Figure 1:**
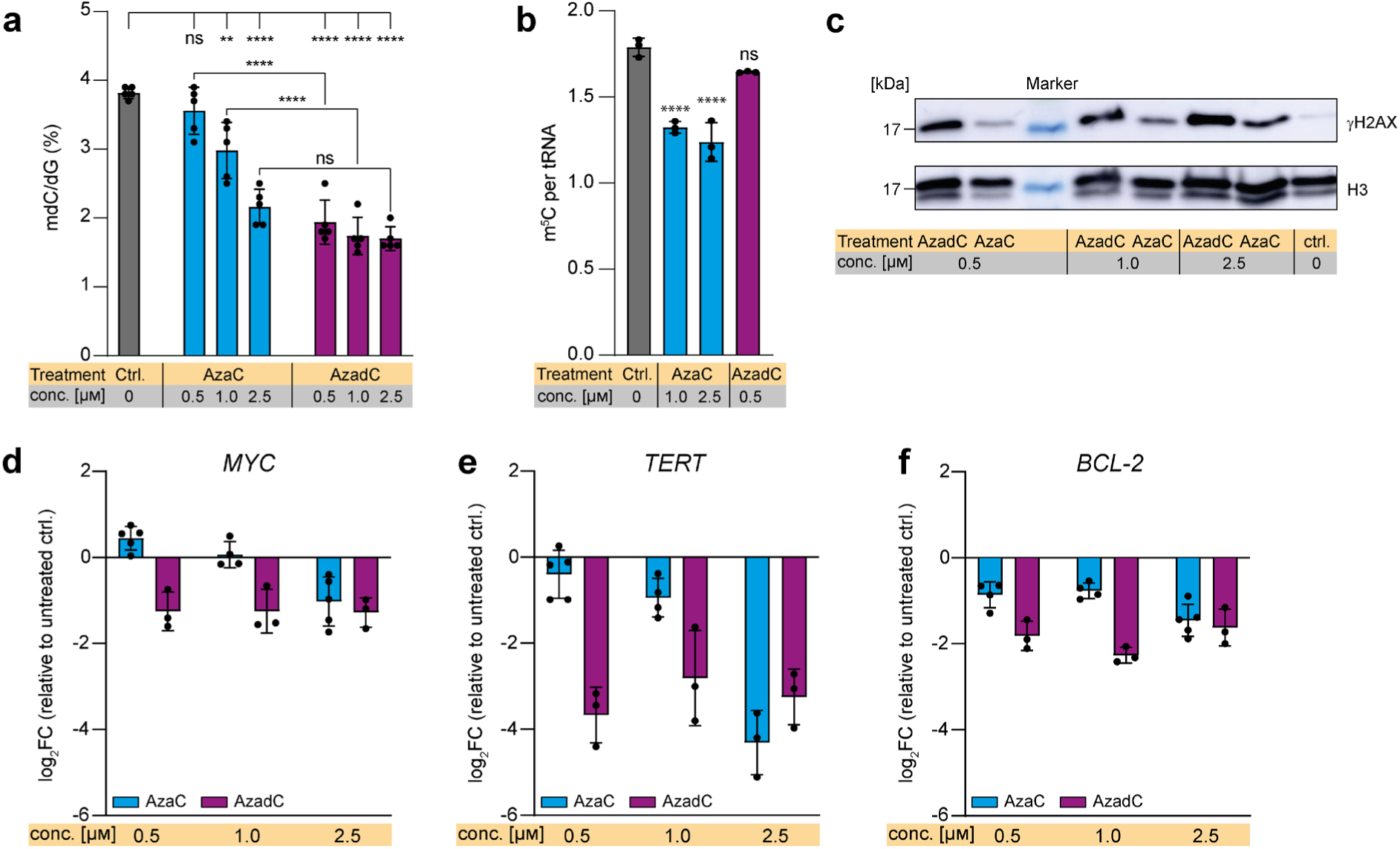
Response of MOLM-13 to AzaC and AzadC treatment on epigenome, DNA damage and transcriptome level. **a** UHPLC-QQQ-MS was used to quantify global mdC levels after exposure to 0.5 µM, 1.0 µM or 2.5 µM AzaC or AzadC for 72 h. Untreated cells served as a control. Ordinary one-way ANOVA with Tukey’s multiple comparisons test was performed. **b** UHPLC-QQQ-MS was used to measure the m^5^C content per tRNA after exposure to 1.0 µM or 2.5 µM AzaC or 0.5 µM AzadC for 72 h. Untreated cells served as a control. Ordinary one-way ANOVA with Dunnett’s multiple comparisons test was performed. **c** Immunoblot analysis of *γ*H2AX levels after exposure to 0.5 µM, 1.0 µM or 2.5 µM AzaC or AzadC for 40 h. Histone H3 served as a loading ctrl. Untreated cells served as a biological control. **d** – **f** RT-qPCR data to quantify gene expression on the transcript level of *MYC* (**d**), *TERT* (**e**) and *BCL-2* (**f**) after 72 h exposure to 0.5 µM, 1.0 µM or 2.5 µM AzaC or AzadC. The log_2_ fold changes (log_2_FC) of the transcripts in relation to the untreated control are displayed. **a** – **b, d** – **f** Each dot represents one independent experiment. Bars show mean, error bars represent standard deviation. All p-values were adjusted for multiple comparisons testing. ns p_adj_ ≥ 0.05, * p_adj_ < 0.05, ** p_adj_ < 0.01, *** p_adj_ < 0.001, **** p_adj_ < 0.0001.

### Transcriptional repression of important oncogenes after AzaC and AzadC treatment

After confirming that treatment with either AzaC or AzadC had the expected effect on MOLM-13 on the epigenetic and DNA damage level, we wanted to further characterize how treatment influenced the phenotype of MOLM-13. Therefore, we investigated how the transcript levels of three important (proto-)oncogenes, namely the transcription factor MYC (Fig. 1d, Figure S1b), the telomerase reverse transcriptase TERT (Fig. 1e, Figure S1c) and the apoptosis inhibitor BCL-2 (Fig. 1f, Figure S1d), all three being main drivers of tumorigenesis and -progression [22], changed upon treatment with AzaC and AzadC. We observed that AzaC was only able to downregulate *MYC* expression at the highest concentration of 2.5 µM (log_2_FC -1.5) and also *TERT* and *BCL-2* expression were only substantially downregulated (log_2_FC *TERT* -3.3, log_2_FC *BCL-2* -1.5) after AzaC treatment at 2.5 µM. In contrast, AzadC treatment was effective in downregulating *MYC, TERT* and *BCL-2* expression at all three concentrations. Importantly, at 2.5 µM, AzaC and AzadC showed a comparable reducing effect on the transcript level of all three genes. These results were in line with the UHPLC-QQQ-MS mdC and the *γ*H2AX western blot data that revealed a similar effect of 2.5 µM AzaC and AzadC treatment.

### Induction of apoptosis with low doses of AzadC, but not AzaC in MOLM-13

Based on these results, a similar effect of AzaC treatment, at least at the highest concentration of 2.5 µM AzaC, and AzadC treatment at all three concentrations on the MOLM-13 cells was expected. Therefore, we monitored the phenotype of MOLM-13 after AzaC and AzadC treatment by brightfield microscopy (Fig. 2a). As an AML cell line, MOLM-13 are suspension cells and show a clearly defined round shape with a diameter of 10 – 15 µm. Surprisingly, the cells mostly contained their defined round shape and cellular diameter after AzaC treatment at all three concentrations, but seemed to have a moderate increase in granularity and proliferated slower compared to the untreated cells. In complete contrast, AzadC treatment at all three concentrations had a severe effect on MOLM-13 morphology and growth. Only a tiny fraction retained the described round shape and anticipated diameter, but instead had dramatically shrunk, indicating advanced apoptosis. A few cells showed enlarged cell size, which is a sign of beginning necrosis (Fig. 2a). When we tested the metabolic activity of MOLM-13 in an MTT assay, which measures metabolic activity to estimate cell viability, proliferation and overall fitness, after treatment with AzaC or AzadC for either 24 h, 48 h or 72 h in a concentration-dependent manner, we also noted substantial differences between the two treatments (Fig. 2b). While the reduction of metabolic activity was only moderate for all tested treatments after 24 h (Fig. S1e), the metabolic activity had decreased significantly in a concentration-dependent manner for all treatments by 48 h. Whereas there was no significant difference between AzaC and AzadC treated cells at the earlier time point, after 48 h AzadC treatment was significantly more effective than AzaC treatment. The average reduction of metabolic activity was between 30 and 40 % for AzaC and between 60 and 70% for AzadC. When exposed for 72 h, no significant additional decrease in metabolic activity in the AzaC treated cells could be observed. However, cells that were treated with AzadC showed an even more drastic drop of metabolic activity of about 80% compared to the control (Fig. 2b). In summary, the MTT assay revealed that AzaC treatment had an initial negative impact on the fitness of the MOLM-13, but metabolic activity could not be reduced further than 50% and only when the highest concentration of 2.5 µM was applied. In contrast, AzadC treatment resulted in a reduction of metabolic activity by 80% for all concentrations tested. Next, we quantified and characterized the type of AzaC-or AzadC caused cell by flow cytometry using a combination of the apoptosis marker Annexin V and a live/dead cell stain (Sytox^™^) (Fig. 2f - h). Annexin^-^/Sytox^-^ cells were considered as viable, Annexin^-^/Sytox^+^ cells as necrotic, Annexin^+^/Sytox^-^ cells as early and Annexin^+^/Sytox^+^ cells as late apoptotic (Fig. 2c). As expected, the majority of the untreated cells was viable with minor populations showing early and late apoptosis and very few undergoing necrosis (Fig. 2c, d). In line with the brightfield microscopy images, AzaC treated cells showed no significant decrease in viability in the flow cytometric analysis. Treatment with the highest concentration of 2.5 µM AzaC did not result in a statistically significant increase of overall apoptotic events compared to the control (Fig. 2d) with cells in the late apoptotic state being absent after 72 h exposure to AzaC at any of the concentrations tested (Fig. 2e). The anticipated increase in granularity from the brightfield microscopy images was, however, not confirmed (Fig. S1f). In contrast, AzadC treatment at all three concentrations resulted in a significant accumulation of cells in an early and late apoptotic, but not in a necrotic state (Fig. 2c – e). Remarkably, the majority of cells was already in a late apoptotic state upon AzadC treatment at only 0.5 µM (Fig. 2c, e). One of the master regulators of apoptosis is BCL-2, which prevents cells from entering apoptosis and is therefore highly expressed in most cancer cells. On transcript level, *BCL-2* was downregulated to the same extent upon application of 2.5 µM AzaC or AzadC, respectively (Fig. 1f), suggesting that apoptosis can be induced after both treatments. However, when analyzing the amount of protein by immunoblot, we observed that 2.5 µM of AzaC failed to downregulate BCL-2 on the protein level in contrast to 2.5 µM AzadC (Fig. 2f, Fig. S2a), indicating that information about the changes on the protein level are essential to judge the effectivity of the treatment. In summary, our data about the cell death events on single-cell level, obtained by flow cytometry, showed that data from MTT or similar assays, which do not determine cell death directly, are not suitable as a stand-alone method to determine viability of cancer cells because the metabolic activity can be influenced by several parameters. In our case, the effects of AzaC on MOLM-13 viability would have been drastically overestimated by MTT assay.

**Figure 2:**
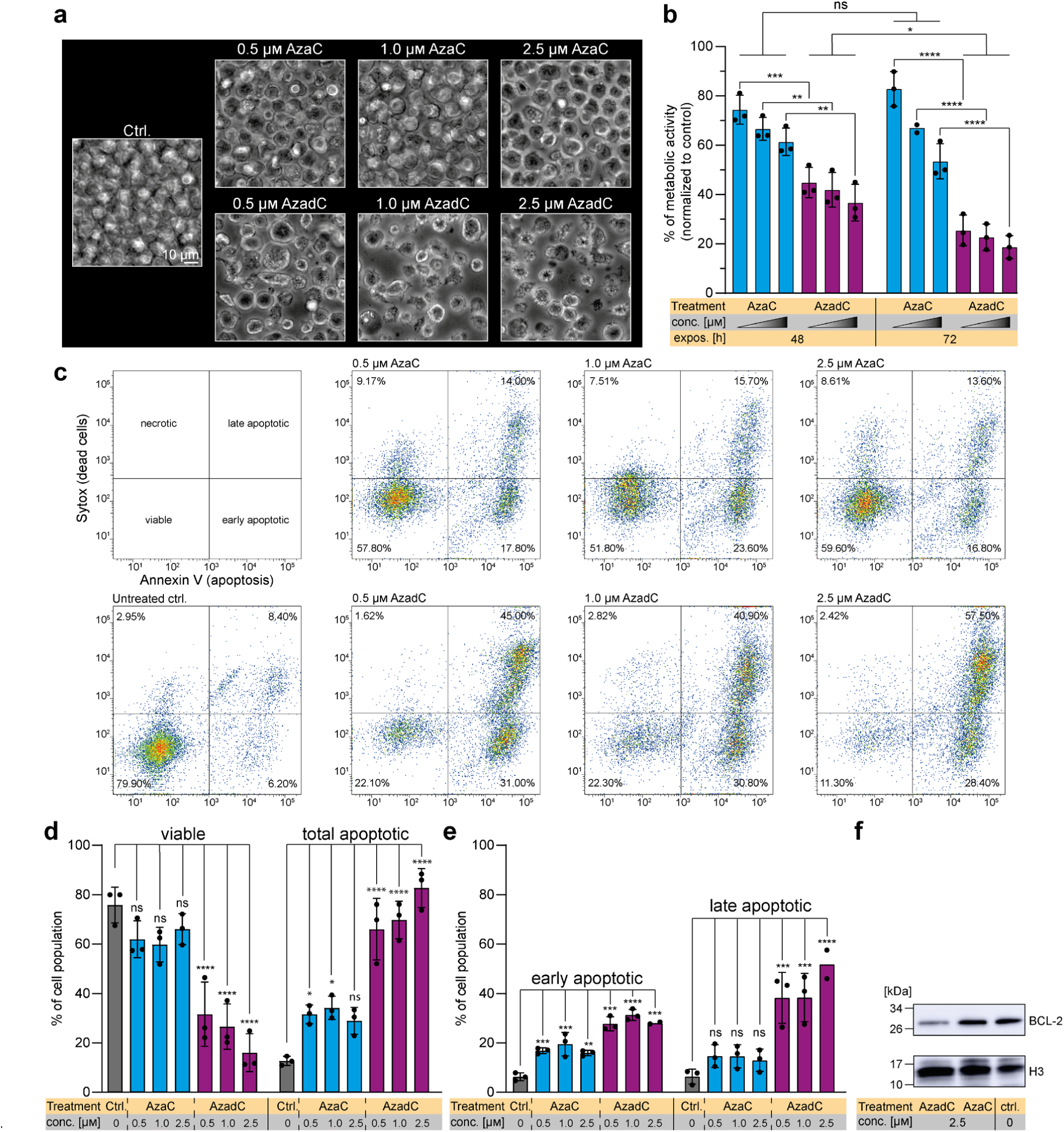
Monitoring of cell death event of MOLM-13 after treatment with AzaC or AzadC. **a** Brightfield microscopy images of MOLM-13 treated for 72 h with different concentrations of either AzaC or AzadC. Untreated cells served as a control. **b** MTT assay results to measure the metabolic activity after 48 h or 72 h exposure to 0.5 µM, 1.0 µM and 2.5 µM (increasing concentrations are indicated by the triangle) AzaC or AzadC. The metabolic activity of treated cells was individually normalized to the metabolic activity of the untreated control for every independent experiment. Ordinary two-way ANOVA combined with Tukey’s multiple comparisons test. **c** Flow cytometric analysis of cell death after AzaC or AzadC treatment in different concentrations. Cells that were Annexin V low and Sytox^™^ low cells were considered viable, cells only high in Sytox as necrotic, cells only high in Annexin V as early apoptotic and cells high in both as late apoptotic. **d** Quantification of three independent flow cytometric cell death analysis performed as depicted in **c** was applied, combining all Annexin V high cells as total apoptotic. **e** Differential analysis of early and late apoptotic events of **d. f** Immunoblot analysis of BCL-2 protein levels upon AzaC and AzadC treatment (2.5 µM each for 72 h) as compared to untreated control. Histone H3 served as a loading control. **b, d, e** Each dot represents one independent experiment. Bars show mean, error bars represent standard deviation. **d, e** Ordinary one-way ANOVA combined with Dunnett’s multiple comparisons test was performed within each observed group (viable and apoptotic in **d**, early and late apoptotic in **e**). **b, d, e** All p-values were adjusted for multiple comparisons testing. ns p_adj_ ≥ 0.05, * p_adj_ < 0.05, ** p_adj_ < 0.01, *** p_adj_ < 0.001, **** p_adj_ < 0.0001.

### Proteome changes after AzaC and AzadC treatment reveal similar, but not identical changes in expression patterns

Since neither the mdC levels, the m^5^C levels, the amount of DNA damage nor the transcript levels of three important oncogenes turned out to be good AzaC or AzadC response markers in MOLM-13 regarding cell death, we aimed to identify other possible cellular markers with better predictive value. To this end, we performed a comprehensive analysis of the proteome changes that were induced by AzaC or AzadC treatment in MOLM-13. We analyzed the proteome changes after 72 h of treatment with drug concentrations that caused a low to moderate induction of cell death, namely 1.0 µM or 2.5 µM AzaC or 0.5 µM AzadC. In total, 129 proteins were differentially expressed compared to the untreated control after exposure to 1.0 µM (Fig. 3a, Table S1) and 161 proteins after exposure to 2.5 µM AzaC (Fig. 3b, Table S2). Regarding the foldchange of all detected proteins, there was a significant positive correlation (Pearson r = 0.6653, Fig. 3c), indicating that AzaC treatment with the two different concentrations had a very similar, but not the same effect on the cells. After 0.5 µM AzadC treatment, 170 proteins were differentially expressed (Fig. 3d, Table S3), among them 95 proteins being downregulated and 75 proteins being upregulated, which was very similar in terms of numbers to 2.5 µM AzaC treatment, where 90 proteins were downregulated and 71 proteins upregulated. Surprisingly, the correlation between the foldchanges of the detected proteins was very similar between 1.0 µM AzaC and 0.5 µM AzadC treatment (Pearson r = 0.6340, Fig. 3e) and 2.5 µM AzaC and 0.5 µM AzadC treatment (Pearson r = 0.5846, Fig. 3f) compared to the correlation between both AzaC treatments (Pearson r = 0.6653, Fig. 3c). Hierarchical clustering revealed that while all four independent replicates of the untreated control clustered together, there was no such clear clustering of the AzaC and AzadC treated cells (Fig. S2b). In line with the flow cytometry data, apoptosis signaling was activated (z-score 1.342, p-value 0.0013) in the pathway analysis for AzadC treatment, but not for AzaC treatment. A closer analysis on the individual differentially expressed proteins revealed that all three treatments (1.0 µM AzaC, 2.5 µM AzaC and 0.5 µM AzadC) resulted in common differential expression of 31 proteins compared to the untreated control (Fig. 3g) and the direction of differential expression (up-or downregulated) was the same. As expected, DNMT1 was significantly downregulated after all three treatments (Fig. 3a, b, d). Myeloperoxidase (MPO), a marker of myeloid lineage commitment, on the other hand was highly upregulated after all three treatments. This is in accordance with literature as the promoter region of MPO is known to be demethylated by DNMT-inhibitors like decitabine, which increases transcription of the *MPO* gene [23]. One of the proteins that showed the highest foldchange in all three data sets was the cellular nucleic acid binding protein (CNBP) (Fig. 3a, b, d). CNBP is a DNA and RNA binding protein that was shown to unfold G-quadruplex structures in promoters of several important oncogenes, thereby promoting their transcription translation on a global scale, among them MYC [24, 25]. Furthermore, two ribosomal proteins, RPL32 and RPL35, were substantially downregulated after all three treatments (Fig. 3a, b, d). We next computed the significance of the overlap of commonly differentially expressed proteins using DynaVenn [26]. The overlap was significant for all treatments and also in a pairwise comparison (p-value < 0.0001). For commonly differentially expressed proteins, the direction of the regulation was always the same. These results indicated that AzaC and AzadC treatment did not result in fundamentally different proteome changes. When analyzing the 93 proteins that were uniquely differentially expressed after 0.5 µM AzadC treatment, there was no enrichment for a specific pathway being altered by AzadC treatment in comparison to AzaC treatment. Rather, AzadC treatment changed the expression of several proteins that are involved in very different cellular pathways, including chromatin organization and heterochromatin formation, transcription and translation, proteasome function and metabolism (Table S3). Interestingly, the deoxynucleoside triphosphate triphosphohydrolase SAMHD1 was among those proteins that were significantly higher expressed after 0.5 µM AzadC treatment only (Fig. 3a, b, d). SAMHD1 is a triphosphate hydrolase that specifically deactivates the AzadC triphosphate, but not the AzaC triphosphate and is therefore one of the reasons why some AML subtypes do not respond to AzadC [27]. However, AzadC treatment resulted in rapid cell death of MOLM-13 despite a fourfold (log_2_FC ∼2) upregulation of SAMHD1, which suggests that the dosage of AzadC must initially be high enough to induce rapid cell death in MOLM-13 before they can efficiently develop resistance mechanisms. In summary, the proteome data showed that AzaC and AzadC treatment had a similar impact on MOLM-13 as indicated by the Pearson correlation coefficients and overlapping differentially expressed proteins, but nevertheless AzaC in contrast to AzadC treatment failed to induce apoptosis effectively at the applied concentrations. This suggests that the accumulative impact of AzadC on various cellular levels was important for cellular fate and not targeting of few defined pathways that were specifically altered by AzadC but not AzaC in MOLM-13.

**Figure 3:**
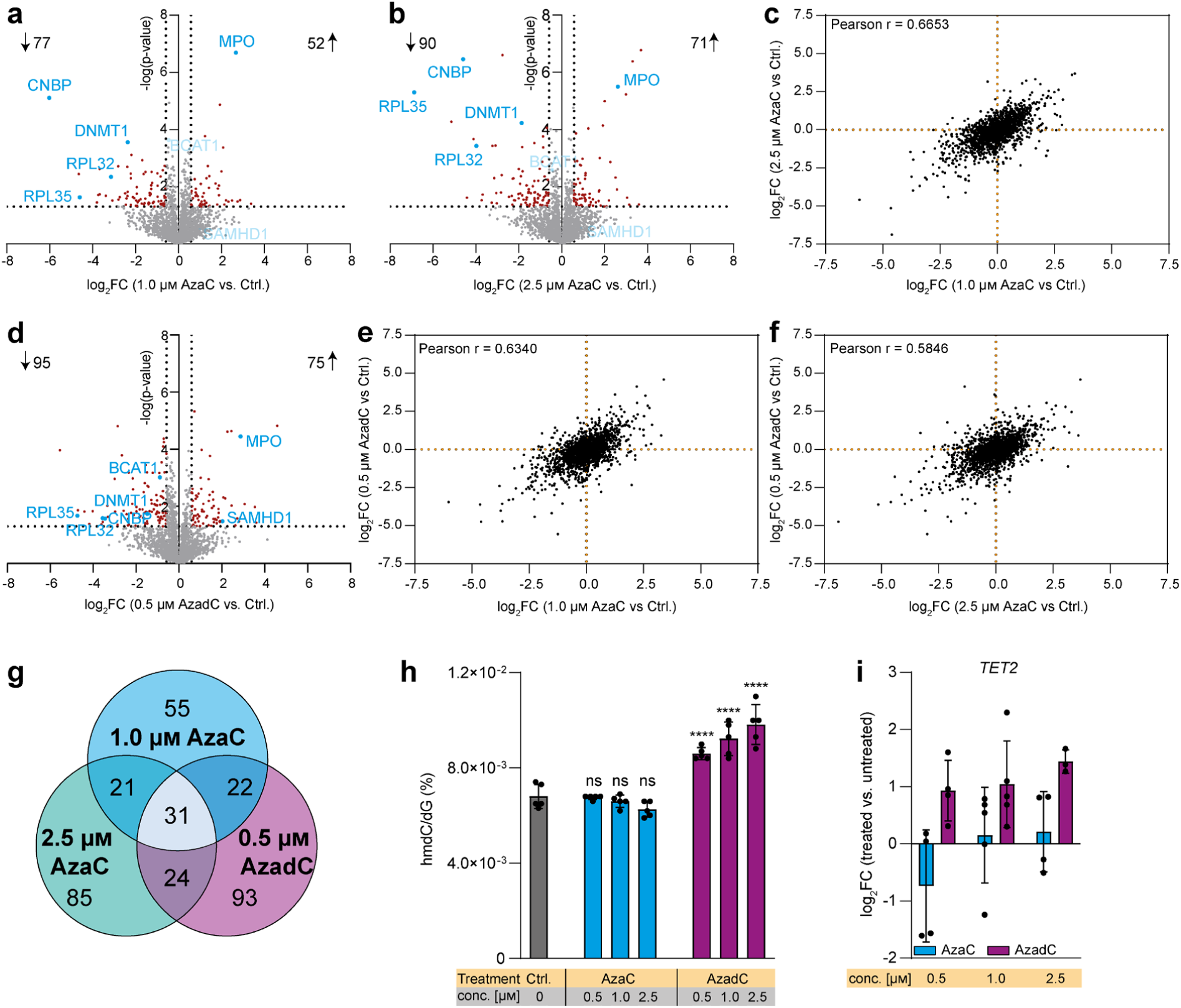
Full-proteome analysis of MOLM-13 and TET activity after 72 h AzaC or AzadC treatment. **a, b** Volcano plot of differentially expressed proteins of **a** 1.0 µM AzaC and **b** 2.5 µM AzaC treated cells versus untreated control. **c** Correlation plot where the log_2_FC after treatment with 1.0 µM AzaC and after 2.5 µM AzaC is displayed for each protein without considering the p-value for the enrichment or depletion. **d** Volcano plot of differentially expressed proteins of 0.5 µM AzadC treated cells versus untreated control. **e** Correlation plot where the log_2_FC after treatment with 1.0 µM AzaC and after 0.5 µM AzadC is displayed for each protein without considering the p-value for the enrichment or depletion. **f** Correlation plot where the log_2_FC after treatment with 2.5 µM AzaC and after 0.5 µM AzadC is displayed for each protein without considering the p-value for the enrichment or depletion. **g** Venn diagram of the differentially expressed proteins after 1.0 µM AzaC, 2.5 µM AzaC and 0.5 µM AzadC treatment. Commonly differentially expressed proteins are indicated by the overlap between two or more treatments. **h** UHPLC-QQQ-MS was used to quantify global hmdC levels after exposure to 0.5 µM, 1.0 µM or 2.5 µM AzaC or AzadC for 72 h. Untreated cells served as a control. Ordinary one-way ANOVA with Tukey’s multiple comparisons test was performed. ns p_adj_ ≥ 0.05, **** p_adj_ < 0.0001. **i** RT-qPCR data to quantify gene expression on the transcript level of *TET2* after 72 h exposure to 0.5 µM, 1.0 µM or 2.5 µM AzaC or AzadC. The log_2_ fold changes (log_2_FC) of the transcripts in relation to the untreated control are displayed. **a, b, d** For each treatment (1.0 µM AzaC, 2.5 µM AzaC, 0.5 µM AzadC, untreated ctrl) four biologically independent experiments were performed and measured. Proteins were considered as differentially expressed when the criteria log_2_FC > |0.58496| (fold change > |1.5|) and -log(p-value) > 1.301 (p-value < 0.05) were both fulfilled. Left side of Volcano plot: proteins depleted after Aza treatment, right side of Volcano plot: proteins enriched after Aza treatment. **c, e, f** log_2_FC for each detected protein of a, b, d, regardless of the p-value, in comparison to each other. Pearson correlation coefficient was calculated. **h, i** Bar shows mean, error bars show standard deviation. Each dot represents one biologically independent experiment.

### hmdC as a possible marker for response to AzaC or AzadC treatment

In the genome, mdC can be further modified by ten-eleven translocation enzymes (TET enzymes), which are *α*-ketoglutarate (*α*KG)-dependent dioxygenases that oxidize mdC to 5-hydroxy-methyl-2’-deoxy-cytidine (hmdC) and further on to 5-formyl-dC and 5-carboxy-dC [28, 29]. It has been shown that cancer cells often have low levels of hmdC and that impaired TET activity, either by reduced TET expression or TET inhibition, promotes leukemogenesis, whereas restored TET activity initiates proliferation stop and differentiation of leukemic cells [30-34]. Therefore, we checked by UHPLC-QQQ-MS how 72 h of AzaC or AzadC treatment changed the global hmdC levels in MOLM-13 and observed that AzaC was not able to increase the hmdC levels at any concentration tested (Fig. 3h). In contrast, AzadC treatment had a concentration-dependent effect. Treatment with 0.5 µM AzadC resulted in a 25% increase, treatment with 1.0 µM AzadC resulted in a 35% increase and treatment with 2.5 µM AzadC resulted in a 45% increase of global hmdC. In contrast to the mdC levels, the hmdC levels therefore correlated with treatment outcome. We further quantified *TET2* transcription levels and observed that AzaC treatment did not lead to increased transcription, but AzadC did (Fig. 3i). Additionally, AzadC, but not AzaC treatment resulted in a significant downregulation of branched-chain aminotransferase 1 (BCAT1) protein (Fig. 3a, b, d). Overexpression of BCAT1 was shown to restrict the *α*KG pool in hematopoietic cells thereby reducing TET activity and promoting leukemic stem cell formation and maintenance [33]. The increased hmdC levels after AzadC treatment could therefore be a consequence of higher TET expression and higher TET activity.

## Discussion

Our data confirm existing reports that mdC and m^5^C levels are not useful biomarkers whether cancer cells respond efficiently to AzaC or AzadC treatment, especially concerning apoptosis induction. Moreover, we could show that even increased formation of DSBs was insufficient to induce rapid cell death in MOLM-13 after AzaC treatment. However, although these markers are not valuable predictors of cell death upon AzaC and AzadC treatment, they can certainly be used to assess whether AzaC and AzadC are taken up by the cells and correctly metabolized, because cellular resistance mechanisms against AzaC or AzadC, e.g. by increased cytidine deaminases or triphosphate hydrolase expression [2], act before the compounds are integrated into DNA and RNA and before DNMT inhibition. The question remains why detrimental effects of DNMT inhibition and thereby formation of severe DNA-protein crosslinks can be tolerated in some cases and not in others. Both compounds induce profound and similar cellular changes on the molecular level in MOLM-13. And yet, the phenotypic outcome was completely different with AzaC failing to induce apoptosis, even at five times higher concentrations compared to AzadC. To understand how the molecular characteristics and the phenotype of cells after AzaC and AzadC treatment are connected, it will be necessary to track AzaC and AzadC metabolism in a holistic approach that uses information from pulse chase experiments and synchronized cells to identify causal time-dependent relationships. In any case it is not sufficient to rely on transcriptome data. Transcriptional downregulation of several key oncogenes with increasing concentration of AzaC resulted in reduced cellular fitness as indicated by the MTT assay but was apparently not sufficient to initiate cell death. Our proteomics data revealed that the translational machinery of MOLM-13 is heavily affected by AzaC but even more by AzadC treatment. Therefore, transcriptome data alone are not enough to estimate the impact of AzaC or AzadC on cancer cells and need to be complemented by proteomics data to gain deeper knowledge about the targeted cellular processes. The only cellular marker in our study that clearly correlated with the failed induction of apoptosis after AzaC treatment was the amount of global hmdC. Interestingly, AzaC treatment did restore *TET2* expression and hmdC in a different leukemic context (T-cell acute lymphoblastic leukemia), which resulted in rapid cell death [35], suggesting that TET activity is an important factor for the efficiency of AzaC or AzadC treatment in inducing cell death. However, whether enhanced TET activity is simply a result of successful cellular reprogramming after AzaC or AzadC treatment or a prerequisite for AzaC or AzadC induced cell death of cancer cells and why AzaC in contrast to AzadC was not able to restore hmdC in the MOLM-13 remains to be investigated. In summary, our study clearly shows that a holistic approach is required to estimate the impact of AzaC and AzadC on cells. It is not sufficient to look at individual aspects like DNA hypomethylation or metabolic activity to predict how well cells respond to AzaC or AzadC treatment. This is especially important to better understand the off-target toxicity of both compounds and to find treatment regimens to avoid them.

## Methods

### Preparation of AzaC and AzadC for cell culture experiments

AzaC (Carbosynth NA02947) and AzadC (Carbosynth NA02969) were purchased and used without further purification. Both compounds were dissolved in Milli-Q H_2_O to a final concentration of 10 mM. Aliquots (10 µL) were made and stored at -80 °C. Directly before usage, the required volume of 10 mM stock solution was thawed and diluted to 100 µM with Milli-Q H_2_O. The integrity of the compounds was checked on a regular basis using HPLC.

### Cell culture MOLM-13

MOLM-13 cells (Leibniz Institute DSMZ-German Collection of Microorganisms and Cell Cultures; AML cell line) were cultivated at 37°C in water saturated, CO_2_-enriched (5%) atmosphere. RPMI 1640 (Sigma-Aldrich R0883), containing 20% (v/v) fetal bovine serum (FBS) (Invitrogen 10500-064) and 1% (v/v) L-alanyl-L-glutamine (Sigma-Aldrich G8541) were used as growing medium. When reaching a density of 2× 10^6^ cells/mL, the cells were routinely passaged to a density of 0.25 – 0.4 × 10^6^ cells/mL. Cells were tested at least once in two months for Mycoplasma contamination using Mycoplasma Detection Kit (Jena Bioscience PP-401L).

### Treatment of MOLM-13 with AzaC or AzadC

For all experiments, cells were seeded at a concentration of 0.5 × 10^6^ cells/mL and directly treated with AzaC and AzadC using the indicated concentration and incubation time during which the medium was not renewed. Untreated cells served as a negative control in all experiments unless indicated otherwise.

### Isolation of gDNA and UHPLC-QQQ-MS

For the determination of mdC levels, 1.5 × 10^6^ cells were seeded and treated with either 0.5 µM, 1.0 µM or 2.5 µM of AzaC or AzadC for 72 h. After 72 h of incubation time, the cells were harvested, gDNA was isolated and UHPLC-QQQ-MS measurements were performed as previously described in *Traube et al*., *2019* [36] with the modification that 3 µg of gDNA per technical replicate were digested using Nucleoside Digestion Mix (NEB M0649S, 2 µL of nuclease, 1.5 h at 37 °C).

### RNA Isolation for RT-qPCR

400 µL of the first flow-through of the gDNA analysis was used to isolate RNA. 300 µL of 95% Ethanol were added, transferred to a Zymo-Spin IIC column (ZymoResearch C1011-50) and incubated for 1 min. After RNA binding, the column was centrifuged at RT, 1500 *x g* (2 min) and at 10 000 x g (30 s). The flow-through was discarded and the column was washed with 800 µL RNA Wash Buffer (ZymoResearch R1003-3). To remove residual DNA contamination, DNA was digested according to the peqGOLD DNase I Digest kit (VWR 13-1091-01), followed by washing steps with 400 µL RNA Prep Buffer (ZymoResearch R1060-2) and 800 µL RNA Wash Buffer. The RNA was eluted using 53 µL of Milli-Q H_2_O and the concentration was determined using a spectrophotometer.

### cDNA synthesis and RT-qPCR

cDNA synthesis was performed with the iScript cDNA Synthesis Kit (Bio-Rad 1708891) according to the manufacturer’s protocol using 1 µg RNA per sample.

For subsequent RT-qPCR the following oligonucleotides were used (Table 1).

**Table 1.**
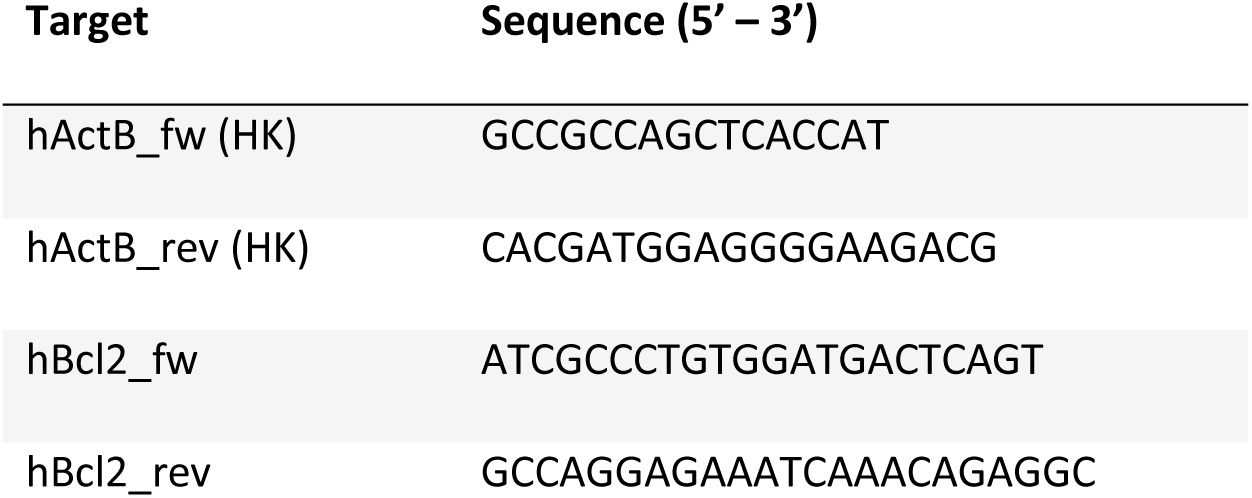
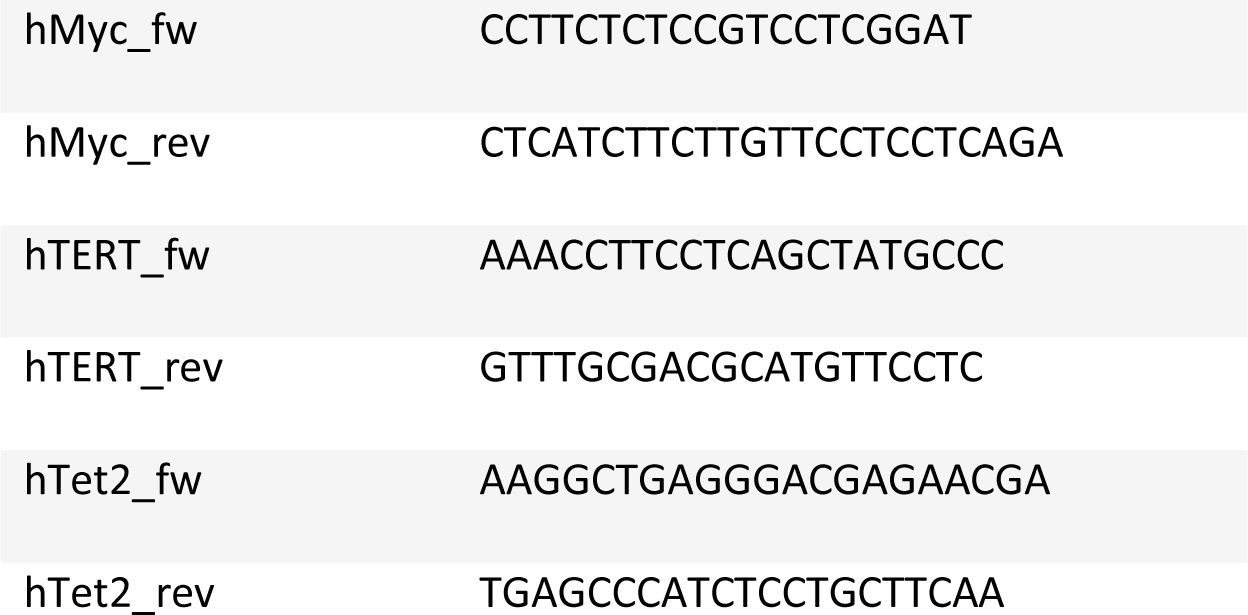
Primers used for RT-qPCR. HK = house keeper

Each primer pair (forward and reverse) was mixed and diluted to a final concentration of 1 µM. Subsequently, per 8 µL of diluted primers, 10 µL of iTaq Universal SYBR Green supermix (Bio-Rad 1725124) was added (RT-qPCR reaction mix). RT-qPCR was performed in 20 µL reactions with 2 µL of 10 ng/µL cDNA (20 ng per reaction) mixed with 18 µL of RT-qPCR reaction mix (final primer concentration 400 nM).

All samples were run in technical triplicates using the following PCR program (Table 2) on a qTOWER^3^/G cycler (Jena Biosciences).

**Table 2.** PCR program used for RT-qPCR

|  |  |  |
| --- | --- | --- |
| 95 °C | 03:00 min. |  |
| 95 °C | 00:10 min. | 40 x |
| 60 °C | 00:20 min. |  |
| 72 °C | 00:30 min. |  |
| Melt | 00:15 min. |  |

An RT–qPCR assay for Actin Beta (ActB) transcripts was used as a housekeeping transcript reference to calculate ΔCt values. Fold change values were calculated with the ΔΔCt method.

### Isolation of RNA for UHPLC-QQQ-MS

For the determination of m^5^C levels, 6.5 × 10^6^ cells were seeded and treated with either 1.0 µM or 2.5 µM of AzaC or 0.5 µM of AzadC for 72 h. After 72 h of incubation time, the cells were harvested, counted and washed once with Dulbecco’s PBS (Sigma-Aldrich D8537-500ML). Afterwards, 1.5 mL of TRI reagent (Sigma-Aldrich T9424-200ML) was added (1 mL for 5 – 10 Mio cells) to lyse the cells. After 5 min incubation time, 150 µL of 1-bromo-3-chloropropane (Sigma-Aldrich B9673) were added (100 µL per 1 mL TRI reagent) and samples were vortexed thoroughly, followed by an incubation time of 10 min. Afterwards cells were centrifuged at 4 °C, 12000 *x g* for 15 min. The resulting red phase contained proteins, the interphase the DNA and the upper transparent phase the RNA. For RNA isolation, the upper phase was transferred into a new 1.5 mL tube and 750 µL of 2-propanol (500 µL per 1 mL of TRI reagent) were added. The samples were vortexed thoroughly and incubated for 10 min at RT, followed by centrifugation at 4 °C, 12000 *x g* for 10 min. The supernatant was discarded and 1.5 mL 75% (v/v) EtOH (1 mL per 1 mL TRI reagent) were added. After having thoroughly vortexed the sample, a centrifugation step at 4 °C, 12000 × g for 5 min was performed. The supernatant was carefully discarded and the pellet was allowed to dry at RT. When the pellet was just about to be completely dry, it was resuspended in 100 µL of Milli-Q H_2_O and the concentration was determined. tRNA was further purified by size exclusion chromatography (SEC) (AdvanceBio SEC 300 Å, 2.7 μm, 7.8 × 300 mm for tRNA combined with BioSEC 1000 Å) according to our previously published protocol [37]. After purification, the RNA was precipitated and dissolved in 30 μL H_2_O. The RNA concentration of each sample was measured using an Implen nanophotometer.

### RNA UHLPC-QQQ-MS

RNA (100 ng) in aqueous digestion mix (15 µL) was digested to single nucleosides and mixed with metabolically produced stable isotope labeled internal standard as previously described [37]. For quantitative mass spectrometry, an Agilent 1290 Infinity II equipped with a diode-array detector (DAD) combined with an Agilent Technologies G6470A Triple Quadrupole system and electrospray ionization (ESI-MS, Agilent Jetstream) was used.

Operating parameters: positive-ion mode, skimmer voltage of 15 V, cell accelerator voltage of 5 V, N_2_ gas temperature of 230 °C and N_2_ gas flow of 6 L/min, sheath gas (N_2_) temperature of 400 °C with a flow of 12 L/min, capillary voltage of 2500 V, nozzle voltage of 0 V and nebulizer at 40 psi. The instrument was operated in dynamic MRM mode (multiple reaction monitoring, MRM). For separation a Core-Shell Technology column (Synergi, 2.5 μm Fusion-RP, 100 Å, 100 × 2 mm column, Phenomenex, Torrance, CA, USA) at 35 °C and a flow rate of 0.35 mL/min were used in combination with a binary mobile phase of 5 mM NH_4_OAc aqueous buffer A, brought to pH 5.6 with glacial acetic acid (65 μL), and an organic buffer B of pure acetonitrile (Roth, LC-MS grade, purity ≥.99.95). The gradient started at 100% solvent A for 1 min, followed by an increase to 10% over 3 min. From 4 to 7 min, solvent B was increased to 40% and was maintained for 1 min before returning to 100% solvent A and a 3 min re-equilibration period. The sample data were analyzed by MassHunter Quantitative Software from Agilent.

### Isolation of nuclear proteins for western blotting

For the preparation of nuclear extracts 2.5 × 10^6^ cells were seeded and treated with 0.5 µM, 1.0 µM or 2.5 µM of AzaC or AzadC for 40 h. After treatment, the cells were harvested and nuclear extracts were prepared as previously described by *Dignam et al*., *1983* [38] with the modification that every buffer was supplemented with Phosphatase Inhibitor Cocktail 2 (Sigma-Aldrich P5726) and Phosphatase Inhibitor Cocktail 3 (Sigma-Aldrich P0044), 1:100 each. Afterwards, the protein concentration was determined using a Bradford assay (Biorad #5000006) as described by the manufacturer. SDS loading buffer (final concentration 50 mM Tris pH 6.8, 100 mM DTT, 2% (w/v) SDS, 10% (v/v) glycerol, 0.1% (w/v) bromophenol blue) was added and the samples were incubated for 5 min at 92 °C before being stored at -20 °C. Before loading the samples on a polyacrylamide gel, the samples were heated for additional 2 min at 92 °C and vortexed thoroughly.

### Isolation of whole proteome for western blotting

For the preparation of nuclear extracts 2.5 × 10^6^ cells were seeded and treated with 0.5 µM, 1.0 µM or 2.5 µM of AzaC or AzadC for 72 h. After treatment, the cells were harvested and washed once with ice-cold PBS. Afterwards, cells were lysed in RIPA buffer (10 mM Tris pH 7.5, 150 mM NaCl, 0.5 mM EDTA, 0.1 % SDS, 2 mM MgCl_2_, 0.5 mM DTT, EDTA-free protease inhibitor cocktail tablet (Roche 43203100), 1 % phosphatase inhibitor cocktail 2 (Sigma P5726-1ML), 1 % phosphatase inhibitor cocktail 3 (Sigma P0044-1ML), DNase I, Benzonase) and incubated for 1.5 h on ice. Afterwards, the lysate was centrifuged at 4 °C, 12000 *x g* for 15 min, the supernatant was transferred into a new tube and and the protein concentration was determined using a Bradford assay (Biorad #5000006) as described by the manufacturer. SDS loading buffer (final concentration 50 mM Tris pH 6.8, 100 mM DTT, 2% (w/v) SDS, 10% (v/v) glycerol, 0.1% (w/v) bromophenol blue) was added and the samples were incubated for 5 min at 92 °C before being stored at -20 °C. Before loading the samples on a polyacrylamide gel, the samples were heated for additional 2 min at 92 °C and vortexed thoroughly.

### Western blotting

15 µg of nuclear extract in SDS loading buffer or 30 µg of total protein extract were loaded on a 4-15% precast polyacrylamide gel (Bio-Rad #4561083EDU) and Color-coded Prestained Protein Marker, Broad Range (10-250 kDa) (New England Biolabs P7719S) was used as a protein standard. The gel was run at constant 150 V for 60 min in SDS running buffer (25 mM Tris, 192 mM glycine, 0.1% (w/v) SDS). For blotting, we used a PVDF blotting membrane (GE Healthcare Amersham Hybond P0.45 PVDG membrane 10600023) and pre-cooled Towbin blotting buffer (25 mM Tris, 192 mM glycine, 20% (v/v) methanol, 0.038% (w/v) SDS). The membrane was activated for 1 min in methanol, washed with Milli-Q water and equilibrated for additional 1 min in Towbin blotting buffer; the Whatman gel blotting papers (Sigma-Aldrich WHA 10426981) were equilibrated for 15 min in Towbin buffer and the precast gel was equilibrated for 5 min in Towbin buffer after the run. Western blotting (tank (wet) electro transfer) was performed at 4°C for 9 h at constant 35 V. After blotting, the PVDF membrane was blocked for 0.5 - 1 h at room temperature using 5% (w/v) milk powder in TBS-T (20 mM Tris pH = 7.5, 150 mM NaCl, 0.1% (v/v) Tween-20). The primary antibodies were diluted in 5 mL of 5% (w/v) milk powder in TBS-T. The blocking suspension was discarded, and the diluted primary antibodies were added for 12 h at 4°C and shaking. After incubation, the primary antibodies were discarded, and the membrane was washed three times ten minutes with TBS-T. HRP-conjugated secondary antibodies were diluted in 5% (w/v) milk powder in TBS-T and added for 1 h at room temperature under shaking. Afterwards, the membrane was washed two times with TBS-T and one time with TBS (TBS-T without Tween-20) before SuperSignal West Pico Chemiluminescent Substrate (Thermo Scientific 34077) was used for imaging. Western blots were imaged using Amersham Imager 680 (auto exposure mode).

For imaging the same blot multiple times using different antibodies, the membrane was directly stripped after imaging. To this end, the membrane was put in TBS-T and the buffer was heated in a microwave until boiling. Afterwards, the buffer was discarded and the procedure was repeated in total three times. After stripping, the membrane was blocked again using 5% (w/v) milk powder in TBS-T and the protocol followed the above described procedure.

Primary antibodies

- Anti-phospho-Histone-H2AX (gH2AX, Ser-139) antibody, Millipore 05-636-1 clone 7BW301, mouse monoclonal antibody, 1:1000
- Anti-Histone-H3 antibody, Cell Signaling Technology 4499S clone D1H2, rabbit monoclonal antibody, 1:1000
- Anti-BCL-2 antibody, ptglab 12789-1-AP Lot 00087156, rabbit polyclonal antibody, 1:1000

Secondary antibodies

- HRP-conjugated anti-mouse IgG, Sigma-Aldrich AP130P, 1:5000
- HRP-conjugated anti-rabbit IgG, Sigma-Aldrich A0545, 1:5000

### Brightfield microscopy

For the brightfield microscopy images, cells were imaged directly in the medium after 72 h treatment with 0.5 µM, 1.0 µM or 2.5 µM of either AzaC or AzadC using an EVOS FL microscope (ThermoFisher) in the transmission mode and 40 x magnification.

### MTT assay

For the MTT assay, 5 × 10^4^ cells were seeded in 100 µL RPMI/20% FBS medium that did not contain phenol red. The assay was performed as described previously[39]. Each timepoint (24 h, 48 h or 72 h) included samples for untreated cells and cells that were treated with 0.5 µM, 1.0 µM or 2.5 µM of either AzaC or AzadC. Each timepoint was measured individually and after measuring the absorption at 570 nm, the average of the technical replicates was calculated and the absorption of the AzaC or AzadC treated cells was set in relation to the absorption measured for the untreated cells, resulting in the relative metabolic activity. Three biologically independent experiments were performed per time point and each sample was measured in technical quadruplicates.

### Flow cytometry analysis

Previous to flow cytometry analysis, cells were treated with 0.5 µM, 1.0 µM or 2.5 µM of either AzaC or AzadC for 72 h, harvested and washed twice with DPBS. Afterwards, cells were counted and 2 × 10^5^ cells per sample were transferred into a new tube. Apoptosis and necrosis were determined by using the FITC Annexin V Apoptosis Detection Kit (BioLegend 640914) and SYTOX^™^ Red Dead Cell Stain (ThermoFisher S34859). To this end, cells were resuspended in 100 µL of Annexin V binding buffer supplemented with 1 µL of FITC-conjugated Annexin V, gently vortexed and incubated at RT for 15 min in the dark. Afterwards, cells were put on ice and 100 µL of cell suspension (2 × 10^5^ cells) were filtered through a 35 µm strainer. 0.2 µL of Sytox were added just before the measurement. For the analysis, BD FACSCanto^™^ and FlowJo Single Cell Analysis Software (v10.8.0) were used. Gates were set once for the control sample and then applied to all other samples.

### Proteomics

MOLM-13 cells were incubated for 72 h with 0.5 μM AzadC, 1 μM or 2.5 μM AzaC in 4 replicates each. Untreated cells (n = 4) served as a control. The cells were harvested and washed twice with PBS. Sample preparation followed in principle a previously published protocol using filter assisted sample preparation (FASP) [40]. Lysis was achieved by adding 1 mL 100 mM TRIS/HCl pH 8.5, 8 M urea, incubation at 95 °C for 5 min and subsequent sonication at 20 % intensity for 20 s using a rod sonicator. The suspension was centrifuged at 14000 × *g* for 5 min and the resulting supernatant was transferred into a new reaction tube. A BCA assay was conducted to determine the protein concentration, before a defined volume (e.g. 700 μL) was taken from each sample and mixed with TCEP (10 mM final concentration) and 2-chloroacetamide (40 mM final concentration) before incubation at 95 °C for 5 min. 30 μg of each sample was filled up to 150 μL with 100 mM TRIS/HCl pH 8.5, 8 M urea and transferred onto a 30 kDa molecular weight cut-off column (Microcon-30, *Merck Millipore*). 150 μL 100 mM TRIS/HCl pH 8.5, 8 M urea was added to the column before centrifugation at 14000 × *g* for 15 min. The flow-through was discarded and the columns were washed twice by addition of 100 μL 50 mM ammonium bicarbonate and subsequent centrifugation at 14000 × *g* for 10 min. The filter units were then placed into a new collection tube before addition of 100 μL 50 mM ammonium bicarbonate containing trypsin (1:100 trypsin:protein) and incubation at 37 °C overnight. Next, the samples were centrifuged at 14000 × *g* for 15 min followed by washing the column twice with 40 μL 50 mM ammonium bicarbonate and centrifugation at 14000 × *g* for 10 min. The eluate was acidified with 5 % formic acid to a pH of 1-2 and purified using C_18_ cartridges (Sep-Pak tC18 1 cc, 50 mg, *Waters*) and a vacuum manifold. For this, the cartridges were washed with 1 mL MeCN, 1 mL 80 % MeCN, 0.5 % formic acid and thrice with 1 mL 0.5 % formic acid, applying vacuum after each step. Then, the acidified samples were loaded onto the cartridges without application of vacuum following three washing steps using 1 mL 0.5 % formic acid. Clean collection tubes were inserted into the vacuum manifold and the peptides were eluted by addition of 250 μL 80 % MeCN, 0.5 % formic acid without applying vacuum before another 250 μL of this buffer was added while applying vacuum. Finally, the solvent was evaporated and the purified peptides were resuspended in 2 % MeCN, 0.1 % formic acid.

2 μg of each sample was submitted to a LC-MS analysis using an *Ultimate 3000 RSLCnano* UHPLC (*Thermo Fisher Scientific*) coupled to an *Orbitrap Eclipse* mass spectrometer (*Thermo Fisher Scientific*) with the *FAIMS Pro* interface attached. First, the peptides were separated by reverse-phase chromatography using a pre-concentration setup. For this, the samples were bound to a precolumn (*Acclaim C18 PepMap100*, 300 μm i.d., 5 mm length; *Thermo Fisher Scientific*) and then eluted onto the analytical column (*SilicaTip Emitter*, 75 µm i.d., 8 µm tip, 15 cm length; *New Objective*; packed in-house with *ReproSil-Pur 120 C18-AQ, 1.9 µm, 120 Å*; *Dr. Maisch GmbH*), which was heated to 40 °C using a column oven by *Sonation*. The separation was achieved at a flow rate of 0.3 μL/min and application of a gradient between solvent A (0.1 % formic acid in H_2_O) and solvent B (0.1 % formic acid in MeCN) going from 7 % to 24.8 % B in 99 min and from 24.8 % to 35.2 % B in 21 min. The ionization was carried out by applying a voltage of 2.0 kV to the column.

The eluting peptides were analysed at alternating FAIMS CV voltages of -50 V and -70 V in *Standard Resolution* mode with the total carrier gas flow set to static and 3.5 L/min. To take peptide charge-state distributions into account, the data dependent acquisition at -50 V was run for 1.7 s, while the corresponding cycle time at -70 V was shortened to 1.3 s. The rest of the parameters were kept identical for the two CV values, starting with a full mass orbitrap scan in profile mode at a resolution of 240000 using a mass range of 375 to 1500 *m/z* and a RF lens level of 30 %. The AGC target was set to *Standard* and the maximum injection time was set to 50 ms. Following this scan, multiple data dependent MS2 scans were carried out for the cycle time specified above using the filters *MIPS* (*Peptide*), *Intensity* (intensity threshold: 1.0e4), *Charge State* (include charge states 2-6, don’t include undetermined charge states) and *Dynamic Exclusion* (exclude for 40 s after one selection with a window of ±10 ppm, also exclude isotopes and other charge states). The most intense ions were individually selected using an isolation window of 1,2 *m/z*, subjected to fragmentation at a normalized HCD energy of 30 % and analysed in the ion trap at *Rapid* scan rate with a *Normal* mass range, the AGC target set to *Standard* and a maximum injection time of 50 ms.

Identification and label-free quantification (LFQ) of peptides and proteins was accomplished using the MaxQuant software version 2.0.3.0 [41, 42]. To prepare the .RAW-files for analysis, they were split into two files containing either the spectra obtained at a CV of -50 V or -70 V. This was accomplished by the *FAIMS MzXML Generator* software developed by the *Coon lab* [43]. During the MaxQuant analysis, these two files were handled like fractions. Next, a database containing the human proteome was chosen. As protease, Trypsin was chosen and a maximum of two missed cleavages were allowed. Carbamidomethylation of C was set as static modification while oxidation of M and phosphorylation of YST was set as dynamic modification. Peptide mass deviations were set to 20 ppm in the first search and 4.5 ppm in the second search, respectively. The minimal peptide length was set to 6 and the PSM and protein FDRs were both set to 0.01. A reverse of the database was chosen as decoy database. The LFQ option was checked and the minimum ratio count was set to 2. The feature “match between runs” was checked and a match time window of 0.7 min as well as an alignment time window of 20 min was defined. Max-Quant results were analyzed using Perseus (v 1.6.14) [44].

### Statistical Analysis

Statistical analysis, except for the proteomics data where Perseus was used, was performed using GraphPad Prism (v 9). Per biological replicate, the mean of the technical replicates was calculated where applicable and only the biological replicates were taken into account for the statistical analysis.

Fig. 1a: Details about the analysis, including exact p-values are given in Table S4.

Fig. 1b: Details about the analysis, including exact p-values are given in Table S5.

Fig. 2b: Details about the analysis, including exact p-values are given in Table S6.

Fig. 2e: Details about the analysis, including exact p-values are given in Table S7.

Fig. 2f: Details about the analysis, including exact p-values are given in Table S8.

## Data Availability

The mass spectrometry proteomics data have been deposited to the ProteomeXchange Consortium via the PRIDE [45, 46] partner repository. The dataset identifier will be added here as soon as available. All other datasets generated for and analyzed in this study are available from the corresponding author on reasonable request.

## Author contributions

TA, CBG and FRT performed cell culture experiments, isolation of gDNA and RNA and mdC/hmdC UHPLC-QQQ measurements. TA and FRT performed flow cytometry measurements. CBG and FRT performed western blots and RT-qPCR. LSR prepared samples for proteomics with help from TA. LSR and FRT analyzed proteomics data. FRT performed MTT assays. NY performed and analyzed m^5^C UHPLC-QQQ measurements under supervision of SK. FRT analyzed and interpreted data, wrote the manuscript with input from all authors and supervised the whole study.

## Acknowledgements

We thank the Deutsche Forschungsgemeinschaft (DFG) for financial support via CRC1309. FRT thanks the Daimler und Benz Stiftung for support. We thank Prof. Thomas Carell (LMU) for having provided the equipment to perform this study and Ria Spallek (TUM) for her input and fruitful discussions.

## Competing interests

The authors declare no competing interest.

## Notes

### Competing Interest Statement

The authors have declared no competing interest.

