## Supplementary Figures for "Comprehensive comparison between azacytidine and decitabine treatment in an acute myeloid leukemia cell line"

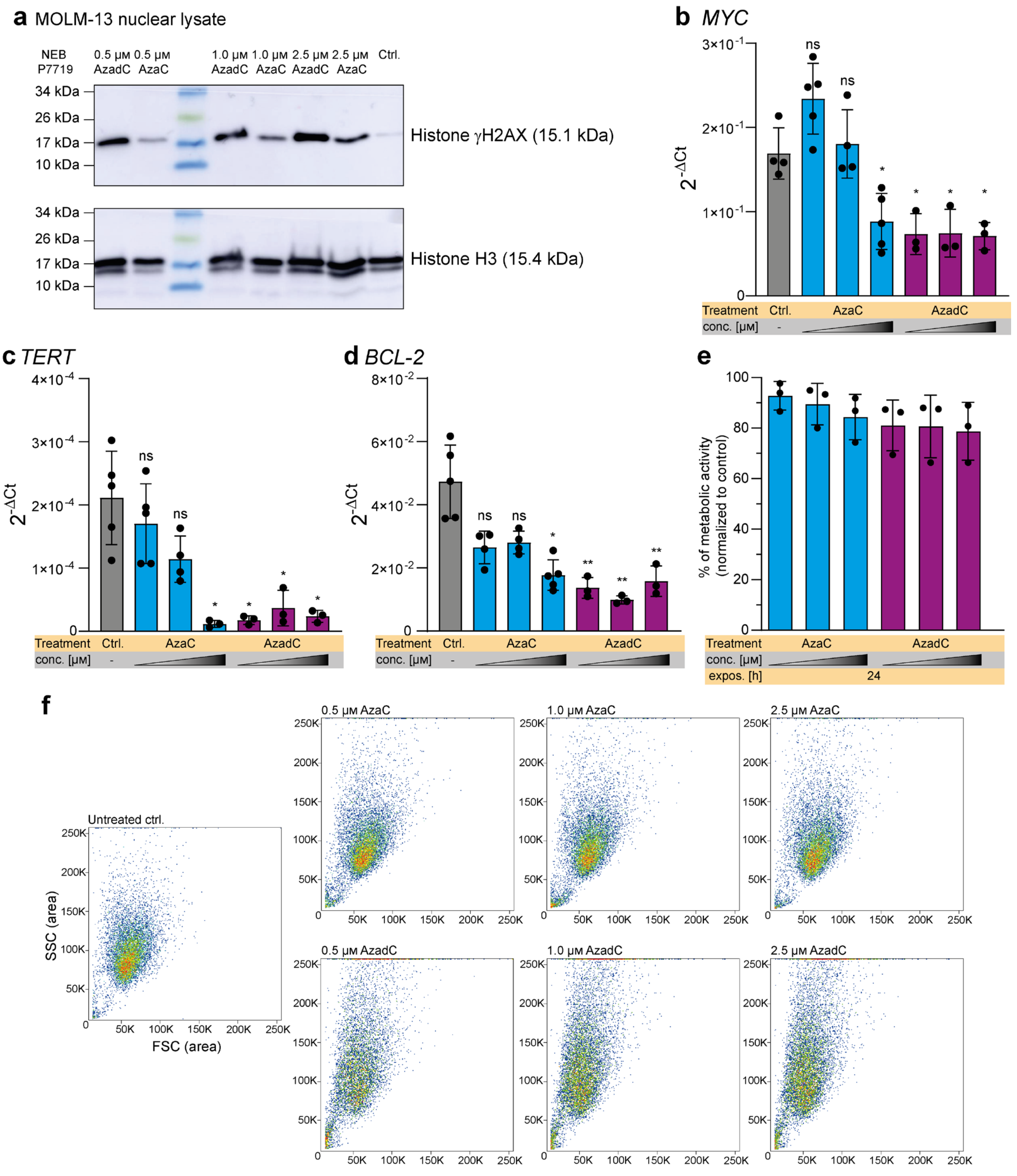


Figure S 1: **Detailed γH2AX immunoblot, $\Delta C_t$ of the RT-qPCRs, MTT assay and flow cytometry**. **a** Detailed immunoblot of Fig. 1c, including the protein ladder. First, γH2AX was detected. Afterwards, the blot was stripped and histone H3 was detected as a loading ctrl. **b** – **d** $2^{-\Delta Ct}$ results of the RT-qPCRs in Fig. 1d – f. Transcripts of interest (TOI) is *MYC* for **b**, *TERT* for **c** and *BCL-2* for **d**, house keeping gene (HK) is Actin **b.** ΔCt = TOI – HK. Statistical analysis Ordinary one-way ANOVA combined with Dunnett's multiple comparisons test. ns $p_{adj} \geq 0.05$; * $p_{adj} < 0.05$, ** $p_{adj} < 0.01$. **e** MTT assay results of MOLM-13 relative to the untreated control after 24 h Flow cytometry analysis of MOLM-13 after increasing concentrations of AzaC or AzadC. Depicted is FSC-A vs SSC-A. **b** – **e** The triangle indicates increasing concentrations of AzaC or AzadC (0.5 µM, 1.0 µM and 2.5 µM).

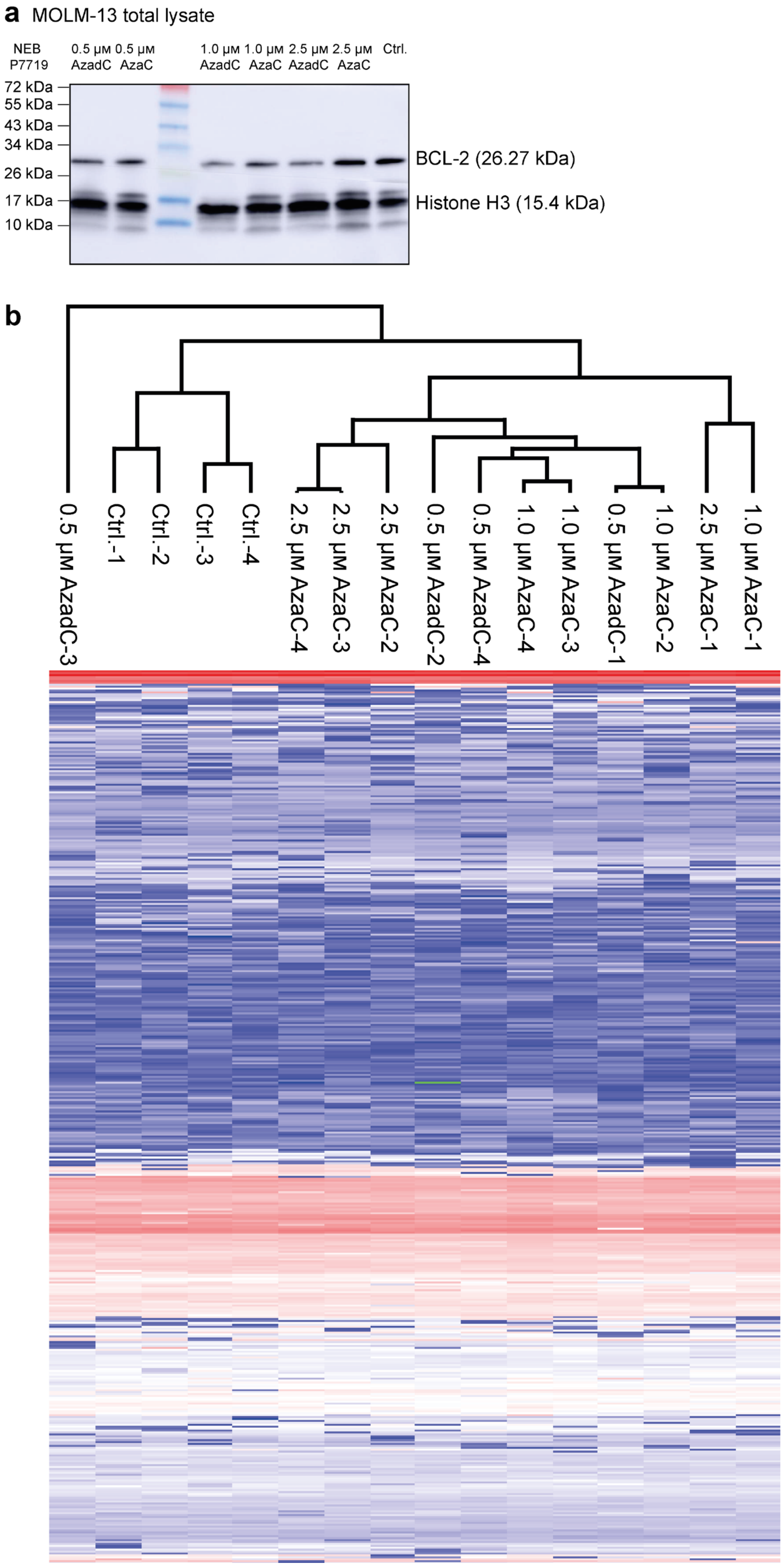


Figure S 2: **Detailed BCL-2 immunoblot and hierarchical clustering of proteomics data sets. a** Detailed immunoblot of Fig. 2f, including the protein ladder. BCL-2 and the loading ctrl. histone H3 were detected in parallel. **b** Hierarchical clustering of proteomics data sets. The heat map was calculated using Euclidian distance for proteomics data sets.
